# Magnetic nanobiopsy: A minimal technology for live cell biomolecular sampling

**DOI:** 10.64898/2026.02.15.705970

**Authors:** Lorena García-Hevia, Débora Muñoz-Guerra, Mónica L Fanarraga

**Affiliations:** The Nanomedicine Group, Instituto de Investigación Valdecilla-IDIVAL, Herrera Oria s/n, 39011 Santander, Spain; Instituto de Investigación en salud Galicia Sur, CINBIO, Universidade de Vigo, 36310, Vigo, Pontevedra, Spain; University of Cantabria, Herrera Oria s/n, 39011 Santander, Spain

**Author notes:** Correspondence: Mónica L Fanarraga, Instituto de Investigación Valdecilla-IDIVAL, Universidad de Cantabria, Herrera Oria s/n, 39011 Santander, Spain.

**Keywords:** Magnetic nanoparticles, Non-destructive cellular sampling, Single-cell heterogeneity, Cellular biomolecular profiling, Longitudinal cell analysis, Precision nanomedicine

## Abstract

Cellular heterogeneity is a fundamental determinant of precision medicine, governing transcriptional plasticity, lineage commitment, and adaptive programs that drive disease progression and therapeutic resistance. However, most molecular interrogation technologies remain inherently destructive, relying on bulk measurements that average cellular signals and systematically obscure rare but functionally decisive subpopulations. As a result, current approaches provide only static snapshots of complex cellular systems, preventing longitudinal analyses of dynamic molecular states and masking the contributions of relapse-initiating, phenotypically plastic, or therapy-resistant cells that ultimately dictate population fate.

Here, we present magnetic nanobiopsy, a simple and scalable platform for minimally invasive, repeatable intracellular molecular sampling from living cells. Magnetically actuated nanocomposites (200 nm–1.4 µm) enable efficient intracellular access and stable biomolecular capture while preserving cellular structural and functional integrity. High-resolution imaging and biochemical analyses confirm robust internalization and anchoring of intracellular biomolecules, while flow cytometry demonstrates that the retrieved cargo remains cell-specific and quantitatively representative of the parental heterogeneous population.

By enabling longitudinal, single-cell–resolved molecular profiling within intact living populations, magnetic nanobiopsy bridges the gap between static bulk analyses and technically complex single-cell methods. This platform establishes a new framework for real-time investigation of cellular heterogeneity, adaptive responses, lineage diversification, and transient cell-state transitions, with broad applicability in cell biology, oncology, and biobanking.

**Highlights:**

- Magnetic nanobiopsy enables non-destructive, longitudinal molecular sampling of living cell populations with high throughput and minimal operational complexity.
- The nanoprobes bypass endo-lysosomal sequestration, directly interfacing with the cytosol to capture representative protein and RNA cargo *via* a membrane-preserving budding exit.
- Quantitative validation confirms that the retrieved molecular profiles faithfully mirror the heterogeneity and relative abundance of the parental cell population.

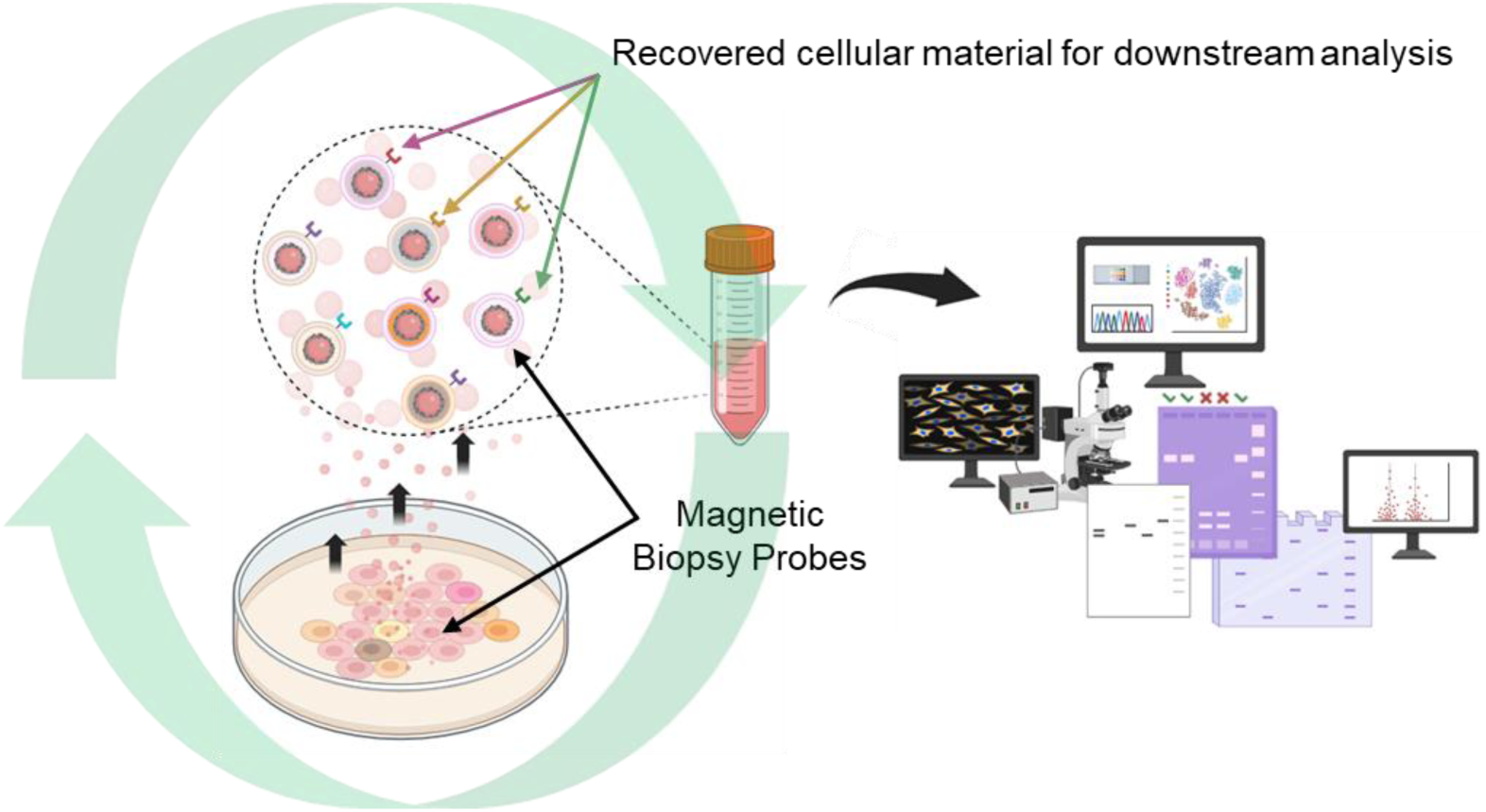

A magnetic nanobiopsy platform is presented for the longitudinal, non-destructive molecular sampling of living cell populations. Magnetically actuated nanocomposites directly access the cytosol—bypassing endo-lysosomal sequestration—to recruit a representative cargo of proteins and RNA through surface interactions and mechanical dragging. Guided by a changing external magnetic field, the molecularly loaded nanoparticles reversibly traverse the plasma membrane and exit via a controlled, budding-like mechanism that ensures cellular integrity. By preserving cell viability and population-level heterogeneity, this approach enables the retrieval of representative biomolecular cargo and the continuous monitoring of dynamic cellular states.

## 1 Introduction

Cellular heterogeneity represents a fundamental barrier to therapeutic precision, frequently acting as the “Achilles’ heel” of clinical intervention by driving treatment failure, disease progression, and unpredictable outcomes across biomedical disciplines. In cancer, even rare drug-tolerant subpopulations can survive initial therapy, persist as quiescent reservoirs, and ultimately fuel recurrence and the expansion of resistant clones [1,2]. Although these phenomena are increasingly recognized, their direct longitudinal interrogation remains technically constrained.

Beyond oncology, intrinsic cellular variability represents a substantial challenge for biobanking and in vitro research. Cell lines and primary cultures—although routinely handled as homogeneous experimental systems—progressively diverge through genetic drift, epigenetic remodeling, chromosomal instability, and microenvironmental selection pressures [3,4]. Continuous cell lines are well known to accumulate karyotypic alterations and subclonal variability over extended passaging, altering transcriptional and functional profiles over time.

Certain populations are particularly susceptible to rapid phenotypic drift during routine expansion, with specialized molecular signatures selectively lost or gained during expansion [5]. For instance, in human mesenchymal cell cultures intended for biomedical applications, molecular drift has been shown to precede cellular senescence, narrowing the window of phenotypic stability and underscoring the need for non-destructive strategies to monitor culture quality longitudinally [6]. Complementarily, Kotrys et al. demonstrated that single-cell fitness and mitochondrial DNA selection are highly dynamic and environment-dependent processes, illustrating how subtle selective pressures can reshape population structure over time [7].

Collectively, these observations indicate that cellular systems are evolving populations rather than static entities. Such instability compromises reproducibility, weakens biological validation, limits the long-term reliability of biobanked resources, and can ultimately undermine therapeutic efficacy. Despite this broad impact, most analytical strategies rely on destructive bulk measurements or technically demanding single-cell methods, restricting repeated intracellular access within the same living population. There is therefore a clear need for simple, non-destructive analytical platforms capable of longitudinal molecular monitoring, enabling direct observation of phenotypic transitions while preserving the biological system under study.

Despite the broad impact of cellular heterogeneity, most analytical strategies remain constrained by either destructive bulk measurements or technically demanding single-cell approaches. Bulk biomolecular analyses average signals from thousands of cells, systematically masking clinically and functionally relevant diversity [8], whereas single-cell technologies—including fluorescence-activated cell sorting (FACS) and single-cell sequencing—have transformed molecular resolution but are inherently destructive [9,10]. By requiring cell isolation and lysis, these methods preclude longitudinal interrogation of the same living population. Consequently, dynamic processes such as the emergence of drug-tolerant states, phenotypic switching, or genomic instability can only be inferred statistically from independently sampled terminal cohorts rather than directly observed.

These limitations underscore the need for minimally invasive platforms capable of repeated, non-destructive intracellular sampling. Longitudinal tracking of molecular trajectories would enable direct monitoring of cell-fate transitions, transcriptional reprogramming, and metabolic adaptation, bridging the gap between static molecular snapshots and the evolving reality of biological systems. Such capability is critical not only for decoding drug-response kinetics in oncology but also for interrogating immune activation, neuronal plasticity, and developmental lineage decisions [2,9–11].

Alternative non-destructive strategies, such as profiling exosomes and extracellular vesicles, offer indirect insight into cellular states but are limited by labor-intensive purification workflows and uncertainty regarding how faithfully they represent the intracellular composition of the parental cell. To address these challenges, a new generation of micro- and nanoscale manipulation technologies—including nanoneedles, atomic force microscopy (AFM) probes, optical and magnetic tweezers, nanowires, and nanoparticle-based systems—has emerged under the concept of “nanobiopsy,” aiming to extract subcellular material while preserving viability [11–15]. A landmark advance was the FluidFM-based nanobiopsy developed by Guillaume-Gentil et al., which enabled controlled cytosolic extraction from single living cells for temporal multi-omic analysis without compromising homeostasis [16]. Similarly, Marcuccio et al. introduced a scanning ion conductance microscopy-based nanobiopsy platform that achieved multigenerational longitudinal transcriptomic profiling of cancer cells, providing high-resolution insights into glioblastoma reprogramming dynamics [17].

While these approaches demonstrate the feasibility and scientific value of longitudinal intracellular sampling, their reliance on specialized instrumentation, high technical complexity, and inherently low throughput—typically limited to individual cells—restrict scalability and application to large, heterogeneous populations. Moreover, sampling a small number of manually selected cells introduces potential stochastic bias and may not capture the functional diversity of the parental system. There is therefore a pressing need for a scalable, high-fidelity intracellular sampling strategy that complements high-resolution single-cell methodologies while delivering the simplicity, robustness, and throughput required to interrogate complex cellular ecosystems without terminal disruption.

Here, we present an innovative nanobiopsy system based on magnetic particles that, guided by an external magnetic field, are controllably introduced and extracted from the cells. This approach enables non-destructive intracellular longitudinal sampling, preserving cell viability.

## 2 Experimental Section

### 2.1 Materials

Fluorescent paramagnetic polystyrene particles of two different types (1% w/v; SPHERO™ Cross-Linked Carboxyl Magnetic Particles, Spherotech Inc.) were employed. These nanocomposites feature an iron oxide layer coated by a polymer layer preventing surface exposure and providing paramagnetic properties. This allows for easy magnetic separation without significant residual magnetism. PS@IO: Fluorescent Sky Blue (Carboxyl, Magnetic) particles, with a diameter of 1.0-1.4 µm (reference FCM-1070-2) and excitation/emission wavelengths of 600/650 nm. PS2@IO: Fluorescent Nile Red (Carboxyl, Magnetic) particles, with a diameter of 0.2-0.39 µm (reference FCM-02556-2) and excitation/emission wavelengths of 500/560 nm. For transmission electron microscopy (TEM) analysis, particles were dispersed in absolute ethanol at a concentration of 0.5 mg/mL. A 10 µL aliquot of the suspension was deposited onto carbon-coated copper grids and air-dried for 1 hour at 25 °C prior to imaging. TEM images were acquired using a JEM-1011 transmission electron microscope (JEOL, Tokyo, Japan) equipped with a high-resolution Gatan digital camera. Particle size distribution was determined by analyzing a minimum of 100 individual particles per sample using ImageJ software (NIH, Bethesda, MD, USA).

### 2.2 Protocol for Magnetic Nanoparticle Cellular Internalization and Biopsy Retrieval

Two different magnetic setups were employed for particle manipulation. For experiments conducted in 24-well plates, a specialized magnetic plate (MF10000; OZ Biosciences, San Diego, CA, USA) containing 24 individual magnets aligned with each well was used [18–21]. For nanoparticle extraction from 60 mm cell culture dishes, a single high-strength neodymium magnet (TCN-63; Webcraft GmbH, Gottmadingen, Germany) was utilized. This magnet has a diameter of 63 mm (±0.1 mm), nickel plating (Ni-Cu-Ni), and a magnetization grade of N38. In both setups, magnetic nanoparticles were introduced into cells by placing the culture plates on the respective magnetic device for 12 hours, allowing cytoplasmic membrane crossing. Particle extraction was performed by repositioning the magnets above the culture plates (lids removed) for 24 to 72 hours to induce nanoparticle exit *via* membrane budding. The released MNPs were then recovered from the culture medium *via* centrifugation.

### 2.3 Cell Culture

HeLa cells (an immortalized human epithelial cell line derived from cervical carcinoma) were cultured in Iscove’s Modified Dulbecco’s Medium (IMDM) supplemented with 10% (v/v) fetal bovine serum (FBS) and 1% penicillin-streptomycin. Cultures were maintained at 37 °C in a humidified atmosphere with 5% CO₂. For magnetic sampling procedures, the cells were incubated with magnetic nanocomposites at a final concentration of 2 µg/mL, resuspended in complete culture medium, for the specified exposure times.

### 2.4 Cell viability and Membrane Integrity Analysis

At the indicated time points, cells were harvested and resuspended in phosphate-buffered saline (PBS). An equal volume of 0.4% trypan blue solution (Thermo Fisher Scientific) was then added to the cell suspension at a 1:1 ratio, and the mixture was incubated for 10 minutes at 37°C. Intact (viable) cells remained unstained, while non-viable cells (those with membrane pores) had their cytoplasm stained blue. Cell counts were performed using a hemocytometer, and viability was calculated as the percentage of unstained cells relative to the total cell population. All measurements were conducted in triplicate to ensure reproducibility.

### 2.5 Cell Imaging

Phase-contrast imaging of live-cell cultures was performed at designated time points using an inverted microscope (Zeiss, Oberkochen, Germany). For fluorescence analysis, cells were imaged under both live and fixed conditions (4% paraformaldehyde). Confocal microscopy was executed on a Nikon A1R system (Tokyo, Japan) equipped with 405, 488, 514, 561, and 638 nm laser lines. All images were processed using NIS-Elements Advanced Research software, with representative fluorescent micrographs presented in pseudo-color. For longitudinal monitoring, time-lapse video microscopy was conducted on cells seeded in Ibidi® μ-dishes (Gräfelfing, Germany). Prior to imaging, cells were incubated with magnetic nanoprobes (MNPs) for 12 h under a static magnetic field applied beneath the culture vessel. Cellular structures were specifically labeled as follows: DNA was stained with 4’,6-diamidino-2-phenylindole (DAPI), RNA was visualized using ethidium bromide (both from Sigma-Aldrich), and cellular membranes were tracked using the lipophilic tracer 3,3’-dioctadecyloxacarbocyanine perchlorate (DiO; Sigma-Aldrich). For ultrastructural analysis, HeLa cell pellets were fixed in 1% glutaraldehyde buffered with 0.12 M phosphate (pH 7.4), post-fixed in 1% osmium tetroxide, and subsequently dehydrated through a graded ethanol series. The samples were embedded in Araldite resin, and ultrathin sections (∼70 nm) were obtained using an ultramicrotome. Sections were contrast-stained with uranyl acetate and lead citrate before being visualized on a JEOL JEM-1011 transmission electron microscope (JEOL, Tokyo, Japan) operating at an acceleration voltage of 80 kV. This high-resolution imaging was employed to characterize the intracellular localization of MNPs and the integrity of the surrounding lipid bilayers.

### 2.6 Genetic Engineering and Fluorescent Protein Expression

To evaluate the extraction of specific intracellular and membrane-bound cargo, HeLa cells were transfected with two distinct genetic constructs: HSP70-eGFP (a gift from Richard Morimoto; Addgene plasmid #15215), serving as a cytoplasmic marker, and a custom TEM8-GFP recombinant construct designed to label the plasma membrane. The latter was engineered by replacing the intracellular domain of TEM8 with a GFP reporter [22]. This synthetic sequence was synthesized by General Biosystems, Inc. (Morrisville, NC, USA) and subcloned into a pcDNA3.1(+) expression vector.

Transient transfections were executed using Lipofectamine™ 2000 (Thermo Fisher Scientific) according to the manufacturer’s protocol. Briefly, 8 µg of plasmid DNA were complexed with the cationic lipid in IMDM and incubated with the cell cultures to achieve optimal expression. Following transfection, cells were treated with magnetic nanocomposites (MNPs) in 5 mL of complete medium for the designated uptake and extraction periods. Finally, the cells were harvested, washed thrice with PBS, and fixed in 4% paraformaldehyde for subsequent fluorescence quantification and imaging analysis.

### 2.7 Flow Cytometry Characterization

To quantify the molecular cargo associated with the extracted probes, both cells and pooled MNPs (recovered via centrifugation) were washed twice with PBS and fixed in 4% paraformaldehyde for 15 min at room temperature. Multiparametric analysis was conducted using a CytoFLEX flow cytometer (Beckman Coulter, Brea, CA, USA) equipped with 405, 488, and 638 nm excitation lasers and thirteen fluorescence detectors. Optical power was standardized at 50 mW for the 488 and 638 nm lines, and 80 mW for the 405 nm line.

To ensure statistical robustness, a minimum of 10,000 events were acquired per sample, with all experimental conditions performed in triplicate. Data acquisition and the subsequent quantification of fluorescent populations (both cellular and MNP-associated) were executed using spectral flow cytometric analysis. This approach enabled precise signal deconvolution and the quantitative assessment of biomolecular anchoring efficiency, providing a high-fidelity comparison between the retrieved material and the parental cell population.

### 2.8 Statistical Analysis

All quantitative experiments were performed with a minimum of three independent replicates. Data are presented as mean ± standard deviation (SD). Statistical comparisons between two groups were performed using a Student’s *t*-test, with statistical significance defined as *p* < 0.05.

### 2.9 SDS-PAGE and Proteomic Profiling

To characterize the protein corona and intracellular cargo recruited by the nanoprobes, whole-cell lysates and the corresponding pooled MNP fractions were subjected to sodium dodecyl sulfate-polyacrylamide gel electrophoresis (SDS-PAGE). Samples were denatured in Laemmli buffer at 95 °C and resolved on 10% polyacrylamide gels under constant voltage. Following electrophoretic separation, total protein content was visualized via staining with Coomassie Brilliant Blue R-250 (Bio-Rad, Hercules, CA, USA). Semiquantitative gel digitization and densitometric analysis were performed using a GelDoc EZ imaging system (Bio-Rad). This comparative proteomic assessment was employed to evaluate the molecular weight distribution of the retrieved proteins and to confirm the qualitative representativeness of the MNP-associated fractions relative to the parental cellular proteome.

## 3 Results

### 3.1 Design and Characterization of Magnetic Nanobiopsy Probes

The development of magnetic nanobiopsy probes (MNPs) is based on the premise that magnetically driven molecular retrieval must occur without compromising cell viability or structural integrity. To address this, we employed two magnetic nanocomposites of different diameters to answer two complementary questions: whether particle size affects the risk of membrane or structural damage, and whether it influences the capacity of the probes to recruit and retrieve biomolecules. Accordingly, we evaluated nanocomposites of 0.2 and 1.4 µm in diameter (Figures 1a, 1b; Experimental Section).

**Fig. 1.**
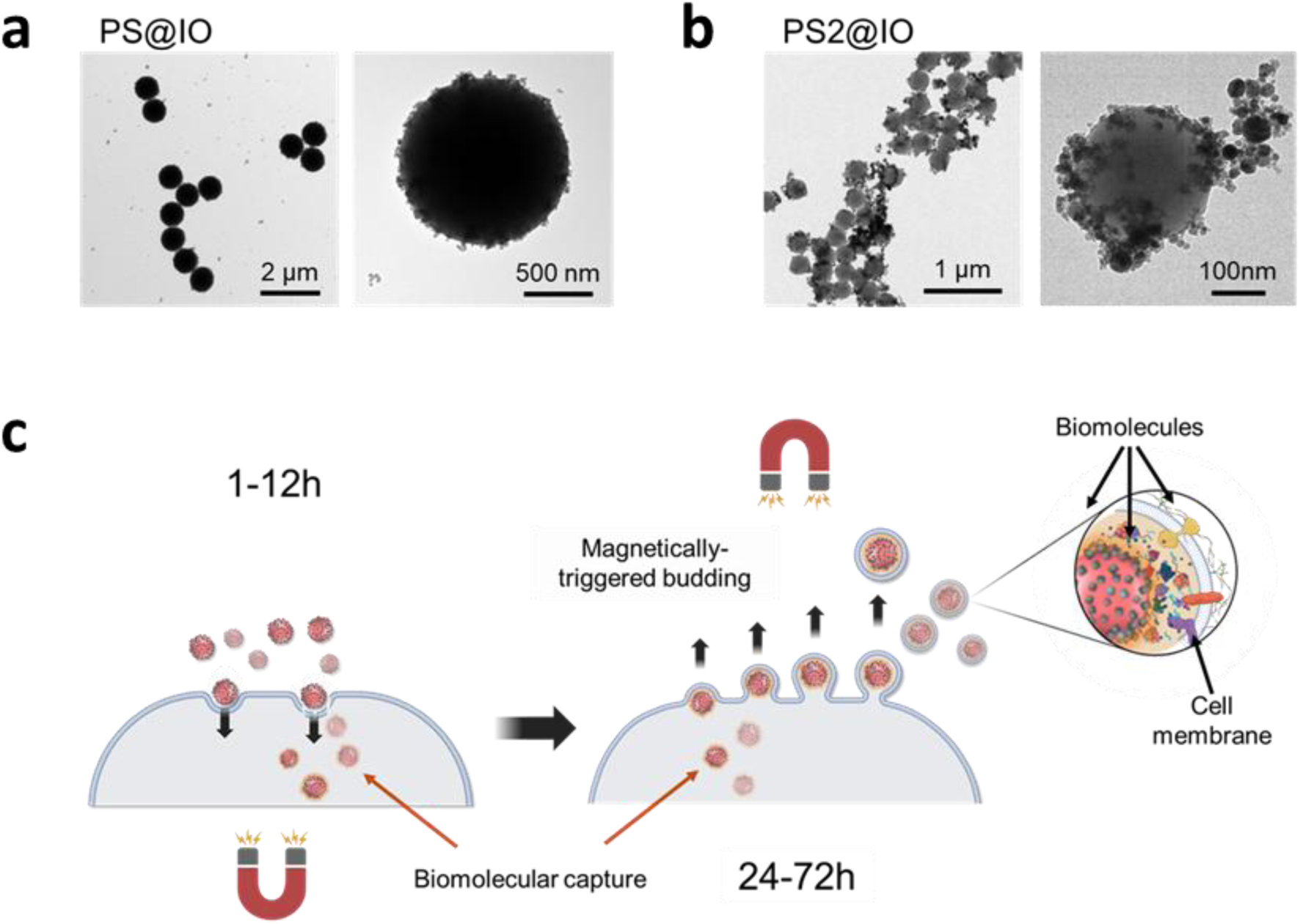
TEM images and schematic of magnetic nanobiopsy. (a–b) TEM images of pristine magnetic nanobiopsy probes: PS@IO (a) and PS2@IO (b), shown at different magnifications. (c) Schematic representation of the magnetic nanobiopsy workflow. During the penetration phase (first 12 h), particles are magnetically driven into the cell by placing a magnet beneath the culture plate. In the budding phase (24-72 h), MBPs are retrieved coated with cytoplasmic and membrane-associated biomolecules, mimicking a viral-like exit mechanism.

Both particle types consist of a fluorescent polystyrene (PS) core uniformly coated with superparamagnetic iron oxide nanoparticles (SPIONs). PS@IO represents the upper size range of the platform (1.0–1.4 µm), whereas PS2@IO provides a smaller, more conservative design (200–400 nm). The intrinsic fluorescence and well-defined size of both nanocomposites enabled accurate detection, intracellular tracking by confocal microscopy, and quantitative analysis by flow cytometry. This dual-size strategy allowed systematic validation of the magnetic nanoprobe (MNP) concept across distinct dimensional regimes and facilitated identification of an optimal balance between magnetic responsiveness, intracellular cargo recruitment efficiency, and biocompatibility.

### 3.2 Maintenance of Cell Homeostasis during Magnetically Induced Entry and Budding

To assess the biocompatibility of the MNPs, they were dispersed in culture medium at a final concentration of 2 µg/mL via bath sonication and immediately added to cultures. Cells were subjected to a standardized two-phase manipulation protocol: (i) *Membrane Penetration Phase* (1–12 h), with the magnet positioned beneath the plate, and (ii) *Budding Phase* (24–72 h), with the magnet repositioned above the plate after lid removal (Figure 1c, Experimental section).

During the membrane penetration phase, the applied magnetic field generated a steep gradient that actively drove nanoparticles toward the cell surface, facilitating direct membrane crossing bypassing all canonical entry methods such as endocytosis. Confocal fluorescence microscopy confirmed intracellular localization of MNPs within the cytoplasm (Figure 2a). These images, complemented by phase-contrast morphological assessment, revealed a normal phenotype consistent across all experimental time points for both types of nanocomposites tested (Figures 2b, S1). The absence of detectable signs of necrosis, apoptosis, or detachment confirms the preservation of cellular homeostasis throughout the protocol, regardless of the nanocomposite composition. Consequently, these results validate the biocompatibility of the magnetically induced entry and budding processes.

**Fig. 2.**
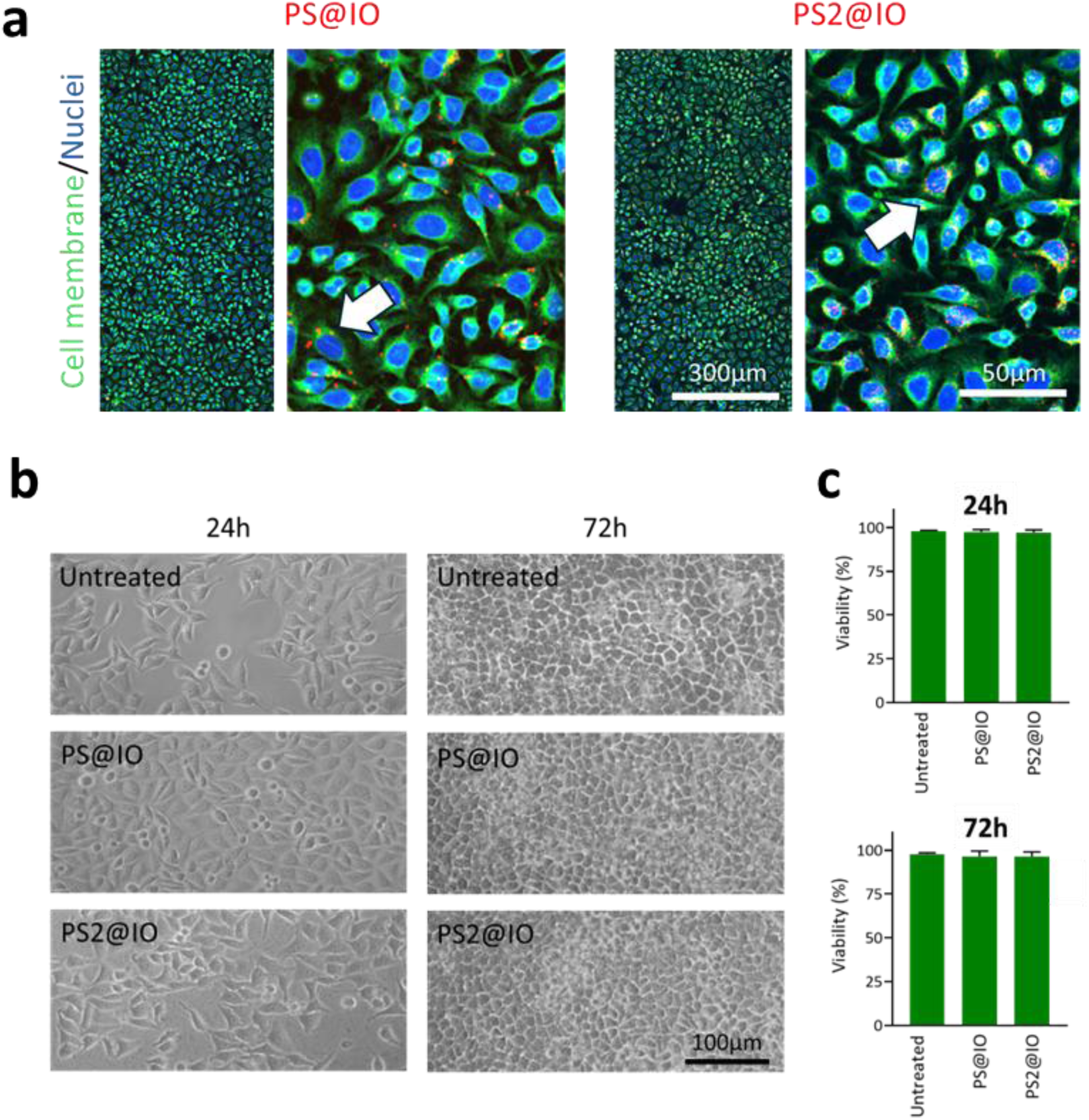
Characterization of cellular interaction and viability during the nanobiopsy process. (a) Representative fluorescence confocal microscopy images of cells interacting with MNPs. Low- and high-magnification views display nuclei (blue), plasma membranes (green), and MNPs (red; white arrows indicate internalized particles). (b) Longitudinal phase-contrast microscopy monitoring of cell cultures during the penetration and extraction (budding) phases compared to untreated controls. (c) Quantitative assessment of cell viability (%) for both MNP-based platforms; no significant deviations were observed compared to untreated cultures (*p* > 0.05), confirming the preservation of cellular homeostasis during magnetic actuation.

Quantitative viability assays confirmed the exceptional biocompatibility of the MNB platform. Across all time points (6, 12, and 24 h), viability rates remained near ∼95%, proving statistically indistinguishable from untreated controls (*p* > 0.05) (Figs. 2c, S2). These data demonstrate that neither the intracellular presence of MNPs nor the magnetic extraction process induces detectable cytotoxicity. Crucially, this non-destructive nature was further validated by sustained cellular health up to 72 h post-sampling, confirming the safety of the budding-mediated mechanism for longitudinal studies.

Both platforms exhibited negligible cytotoxicity, with no detectable signs of necrosis or apoptosis. Consistent with previous studies employing AFM probes or scanning ion conductance microscopy (SICM) for membrane penetration [16,17], our findings confirm that the transient physical perforation of the lipid bilayer during particle entry does not compromise cellular survival. Furthermore, the retrieval phase mimics physiological pathways such as viral egress or extracellular vesicle shedding. As a biomimetic and inherently less disruptive process than mechanical puncture, this induced budding-off ensures the maintenance of membrane continuity. Consequently, these observations validate the magnetic nanobiopsy platform as a minimally invasive and biocompatible method for longitudinal molecular sampling.

### 3.3 Intracellular Penetration, Localization, and Biomolecular Anchoring

Having established that MNP translocation does not compromise cellular homeostasis, we next characterized their intracellular fate and capacity to interact with and recruit cytosolic components. Specifically, we examined the subcellular distribution of the MNPs and the nature of their interactions with the intracellular environment during their cytoplasmic residence. Confocal microscopy at 24 and 48 h confirmed that cells maintained a healthy-looking morphology, with no detectable signs of apoptosis, necrosis, or cytoplasmic stress - even in cells harboring multiple probes. MNPs were predominantly localized within the cytoplasm, exhibiting a marked perinuclear enrichment (Figures 3a, S3 and S4). Critically, the microtubule network remained intact with no observable disruption of the cytoskeletal architecture, further substantiating the excellent cytocompatibility of the nanoprobes.

**Fig. 3.**
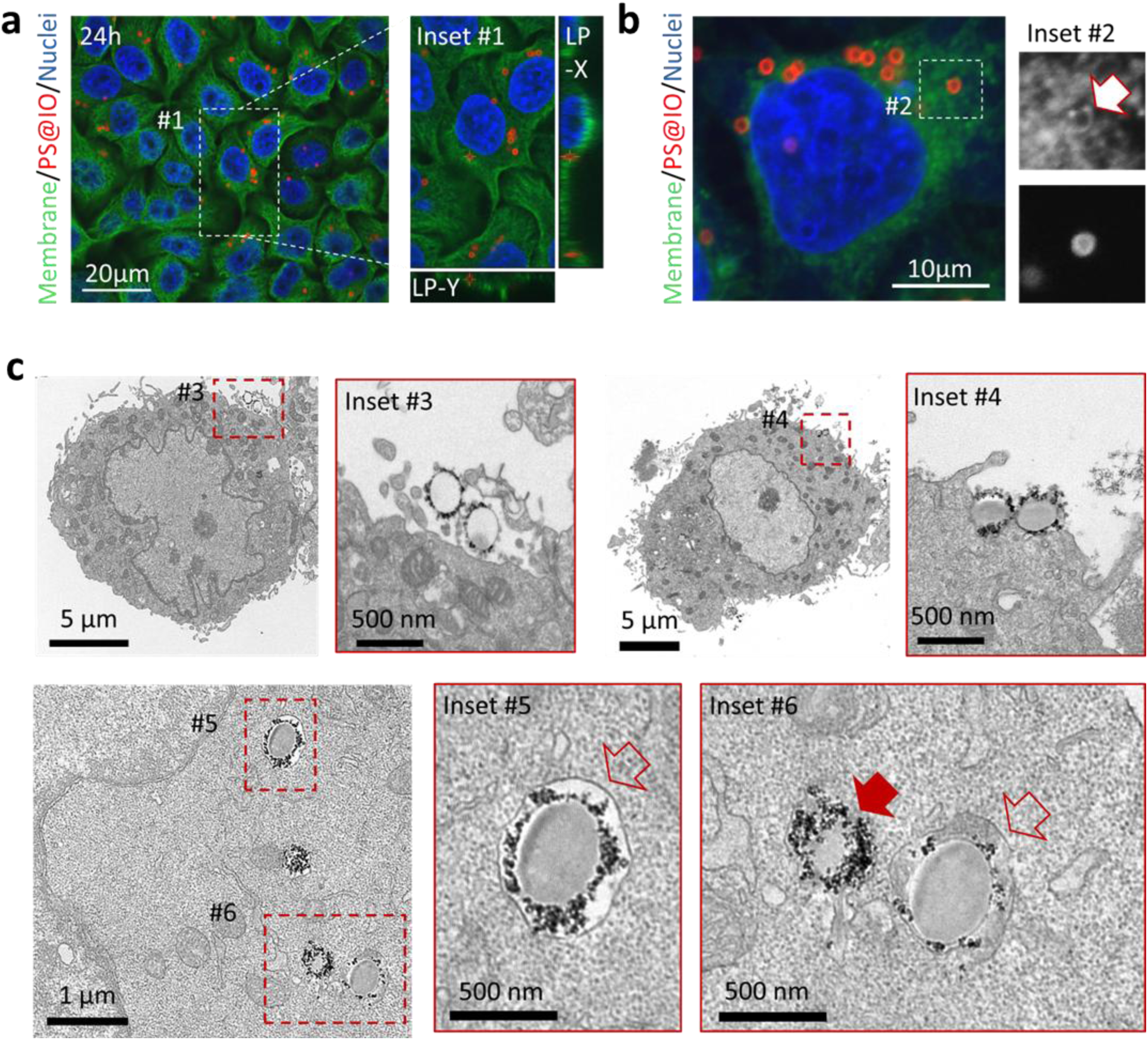
Intracellular translocation and biomolecular recruitment. (a) Confocal microscopy images of cells after 24 h of incubation with PS@IO nanoparticles (red); plasma membranes are stained with DiO (green). Inset 1 shows orthogonal (XZ) projections confirming the internal localization of the MNPs. (b) High-resolution single Z-plane confocal image of an individual cell harboring multiple MNPs. Inset 2 reveals a green fluorescent halo (white arrow) enveloping a particle, indicating the acquisition of a membranous coating. (c) TEM micrographs of ultrathin sections providing ultrastructural evidence of the nanobiopsy process. Insets 3 and 4 capture MNPs in the early stages of interaction, from surface contact to active plasma membrane translocation. Inset 5 displays an internalized MNP sequestered within an intact vesicular structure (red open arrow). Inset 6 highlights the diversity of intracellular interactions: MNPs freely intermingling with the cytosol and ribosomes (red solid arrow), and a particle associated with local membranous fragments (red open arrow).

High-resolution single Z-plane confocal imaging demonstrated that internalized MNPs were closely associated with intracellular structures. In representative examples, individual probes were enveloped by membrane-derived components, forming a characteristic ring-like configuration (Figure 3b, inset 2, arrow). This suggests an active engagement with intracellular membranes, potentially facilitating the capture of a representative biomolecular coating prior to retrieval. While the majority of the probes remained cytosolic, occasional particles were detected within the nuclear compartment (Figure S4, arrow), as confirmed by orthogonal projections.

Whole-cell fluorescence observations were corroborated by ultrastructural analysis via TEM. Imaging of ultrathin sections confirmed the presence of MNPs at discrete stages of the magnetic manipulation process, specifically during the transmembrane entry and budding-mediated exit steps. Particles were captured in direct contact with or actively traversing the plasma membrane (Figure 3c, insets 3 and 4), providing high-resolution evidence of efficient membrane translocation under magnetic actuation.

Within the intracellular environment, MNPs were distributed throughout the cytoplasm, frequently localized in the perinuclear region (Figure 3c, inset 5). At this ultrastructural level, distinct interaction patterns were observed: some particles appeared fully sequestered within membranous compartments, while others were integrated into the cytosol, closely surrounded by ribosomes or vesicular structures (Figure 3c, inset 6, arrows).

Importantly, confirming our previous light microscopy observations, no significant ultrastructural alterations were detected. Organelles, endomembrane systems, and the overall cellular architecture remained pristine, indicating that the mechanical stress associated with magnetic manipulation does not induce detectable structural damage. This preserved ultrastructure validates the feasibility of longitudinal or sequential nanobiopsy sampling from the same living cell without compromising its biological integrity.

### 3.4 Biomolecular Recruitment and Mechanical Dragging

To assess the capacity of the MNPs to act as molecular carriers, we first examined fixed cells 24 h post-incubation using ethidium bromide to visualize intracellular nucleic acids. Confocal imaging revealed a robust recruitment of nucleic acids onto the MNP surface. High-resolution single Z-plane imaging displayed a distinct, doughnut-shaped fluorescent halo surrounding the particles, consistent with the adsorption of cytoplasmic nucleic acids—predominantly RNA—forming a stable biomolecular corona (Figure 4a, inset 1).

**Fig. 4.**
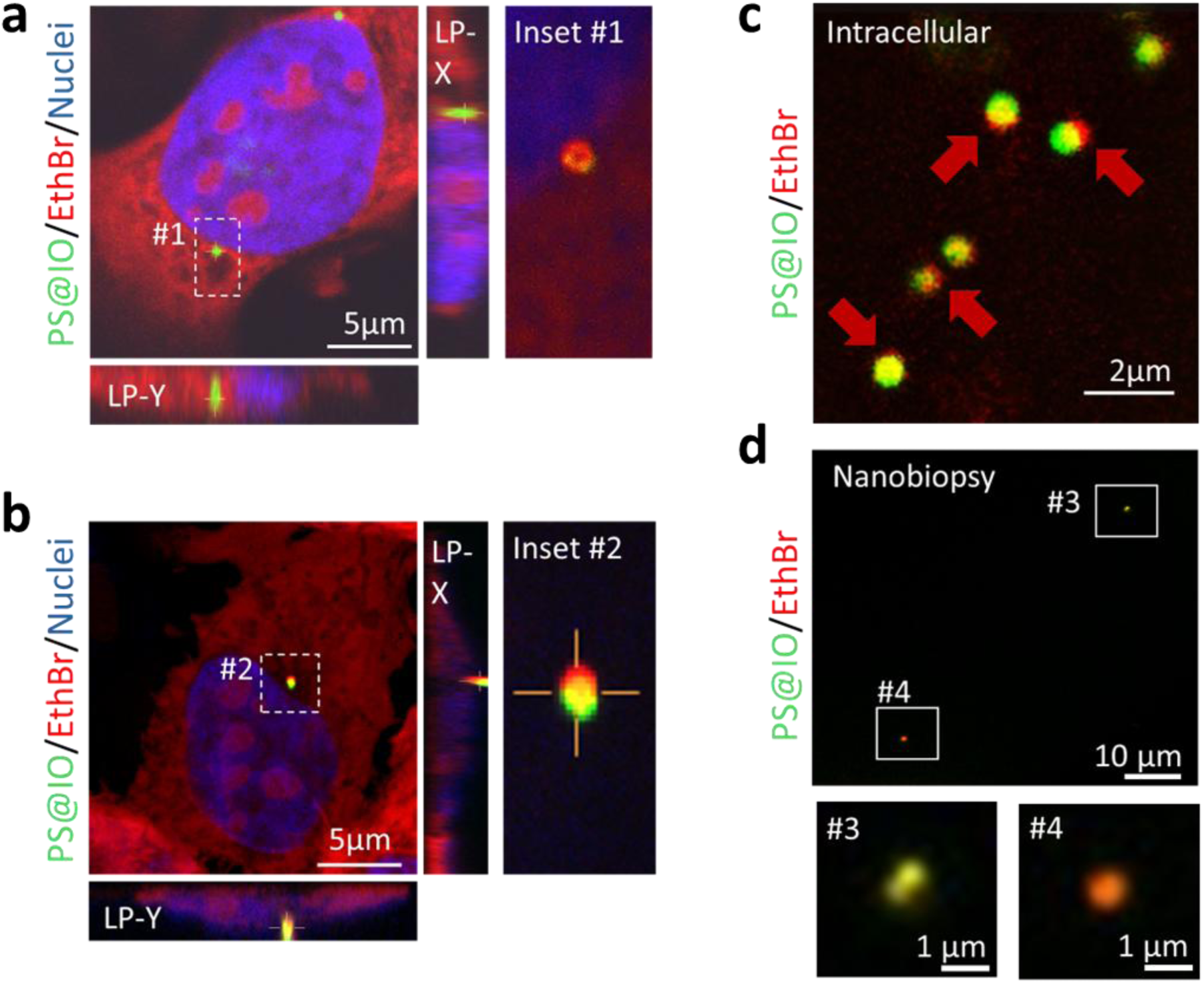
Biomolecular recruitment, intracellular dragging, and successful cargo retrieval. (a) Confocal projection and orthogonal (XZ) views of a cell containing a single PS@IO nanoparticle (green), confirming intracellular localization. Nucleic acids are stained with ethidium bromide (red). Inset 1 (single Z-plane) reveals a dense, doughnut-shaped nucleic acid halo surrounding the MNP. (b) Single Z-plane and orthogonal (XZ) projections showing an MNP trailing a comet-like nucleic acid pattern; Inset 2 provides a high-resolution view of the tail. (c) Live-cell time-lapse frame (Video S1) demonstrating dynamic dragging of stained nucleic acids by moving MNPs (arrows). (d) Confocal analysis of retrieved MNPs confirming successful capture and retention of nucleic acids after extraction in a coated MNP (Insets 3 and 4).

Interestingly, several intracellular MNPs were associated with “comet-tail–like” trails of stained material (Figure 4b). This initial observation in fixed samples suggested that nucleic acids remained firmly attached to the MNP surface, forming these structures as the particles were displaced through the crowded intracellular environment. To validate this phenomenon in a physiological context, we performed high-resolution time-lapse confocal microscopy on living cells. This dynamic analysis unequivocally confirmed the existence of these nucleic acid tails (Figure 4c and Video S1). As the MNPs moved through the cytoplasm, they generated clearly visible trails of stained genetic material, providing direct evidence of mechanical dragging and stable biomolecular sequestration. These results confirm that the MNP surface acts as a functional scaffold capable of retaining a representative molecular cargo during its intracellular journey. Finally, confocal analysis of the retrieved probes revealed a distinct fluorescent signal from the ethidium bromide-stained cargo, mirroring the patterns observed intracellularly (Figure 4d). This confirms that the nucleic acids remain successfully anchored to the MNP surface throughout the magnetic extraction and recovery process.

Together, confocal and TEM analyses demonstrate efficient cytoplasmic access, intracellular molecular anchoring, and biocompatible retrieval of MNPs. These findings confirm that the probes bypass endo-lysosomal sequestration, directly interface with cytoplasmic biomolecules, and function as non-destructive tools for minimally invasive, longitudinal molecular sampling in living cells.

### 3.5 Retrieval and Characterization of the Molecularly Anchored Cargo

Following magnetic extraction, the molecularly loaded MNPs were recovered from the culture medium and subjected to ultrastructural and biochemical characterization. TEM imaging provided direct confirmation of biomolecule acquisition; extracted MNPs were frequently enveloped by an irregular membranous coating, consistent with the vesicle-mediated budding observed during the exit step (Figure 5a). High-resolution sections revealed the characteristic architecture of the retrieved probes: a polystyrene core coated with iron oxide nanoparticles, now overlaid with a distinct layer of cell-derived membranous and biomolecular material.

**Fig. 5.**
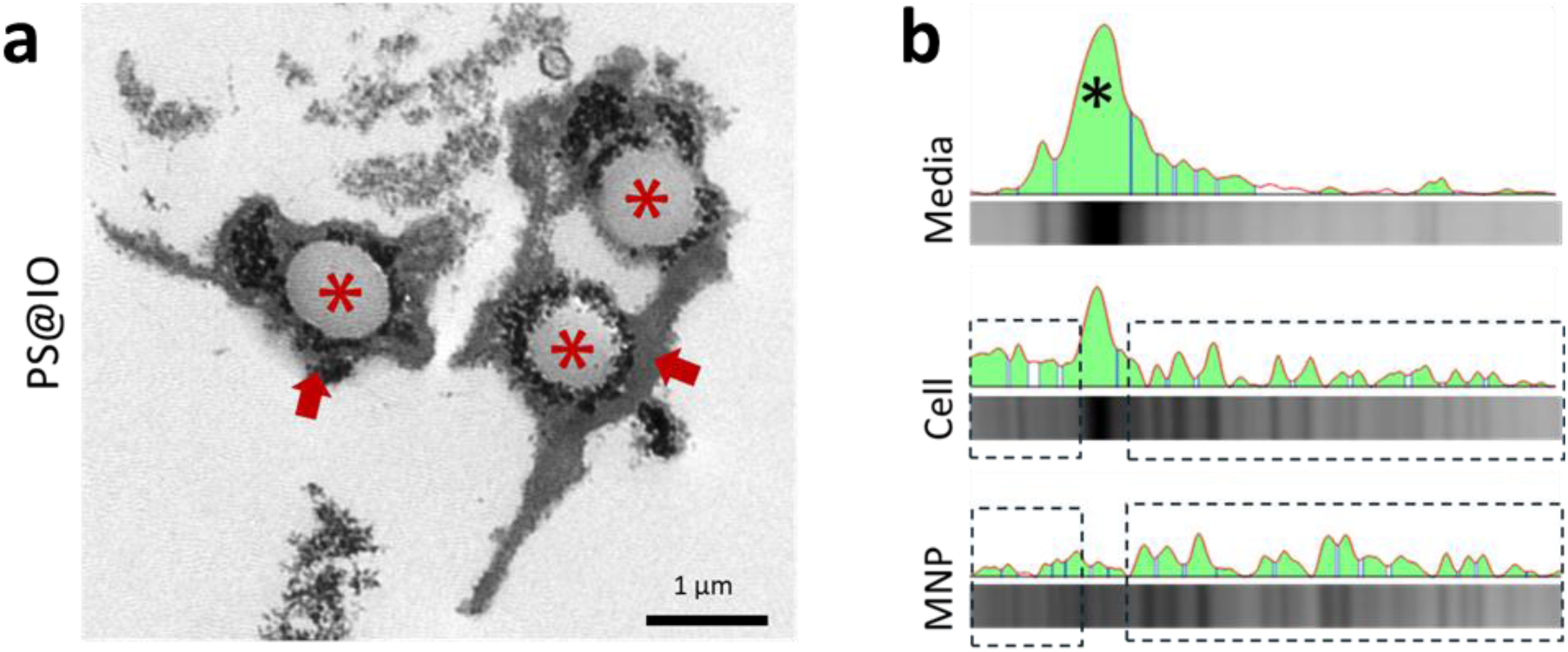
Ultrastructural and biochemical validation of retrieved cargo. (a) TEM micrograph of an ultrathin section showing three MNPs extracted from cells. The polystyrene core (red asterisk), the surrounding magnetic nanoparticle layer, and the associated biomolecular coating are clearly distinguishable (arrows). (b) Qualitative and semi-quantitative densitometric profiles (green) derived from SDS–PAGE (Coomassie-stained) of proteins recruited from the culture medium, parental cell lysate, and the pooled MNP fraction. The banding pattern and protein landscape of the MNP fraction show a high degree of correlation with the parental cell lysate (boxes), confirming the selective retrieval of intracellular components over non-specific adsorption from the culture medium. The most abundant protein in serum, albumin, is indicated with a black asterisk.

To validate the cellular origin of this cargo, isolated MNPs were pooled and analyzed via SDS–PAGE. The protein profile of the retrieved MNPs was compared against both the whole-cell lysate and the serum-containing culture medium. Coomassie-stained gels revealed multiple protein bands associated with the nanobiopsy samples (Figure S5). Crucially, the protein landscape of the retrieved MNPs was clearly distinct from the serum components of the culture medium, while closely mirroring the distribution pattern of the parental cell lysates (Figure 5b). This indicates a selective enrichment of intracellular proteins rather than nonspecific adsorption from the extracellular environment.

Collectively, these analyses demonstrate that MNPs effectively capture and transport intracellular biomolecules. The recovered cargo is compatible with downstream proteomic and transcriptomic workflows, validating the magnetic nanobiopsy as a robust, minimally invasive strategy for molecular sampling from living cells.

### 3.6 Quantitative Valuation of Cellular Diversity via Nanobiopsy

To evaluate if magnetic nanobiopsy accurately reflects cellular heterogeneity, we sampled both the cytoplasm and plasma membrane—environments of high translational value for retrieving stress-related regulatory proteins and pathological surface signatures, respectively. This dual-reporter approach allowed us to assess the MNBs’ capacity to recruit cargo from both fluid cytosolic and structurally anchored membrane environments, determining whether the ensemble of recovered probes effectively mirrors the molecular abundance of the parental population.

As a technical demonstration, cells were transfected with either Hsp70:GFP (cytoplasmic protein) or TEM8:GFP (membrane-bound receptor). The inherent fluorescence of these reporters served as a straightforward analytical readout, enabling a straightforward and robust quantification of protein presence across both cells and probes. Confocal microscopy and whole-cell flow cytometry initially characterized the parental populations (Figs. 6a–d), showing expression levels of approximately 60–70%. To determine if the retrieved samples were representative of these populations, we analyzed the extracted MNPs using flow cytometry (Fig. S6), quantifying the fraction of fluorescently loaded probes relative to the original culture (Figs. 6e, f).

**Fig. 6.**
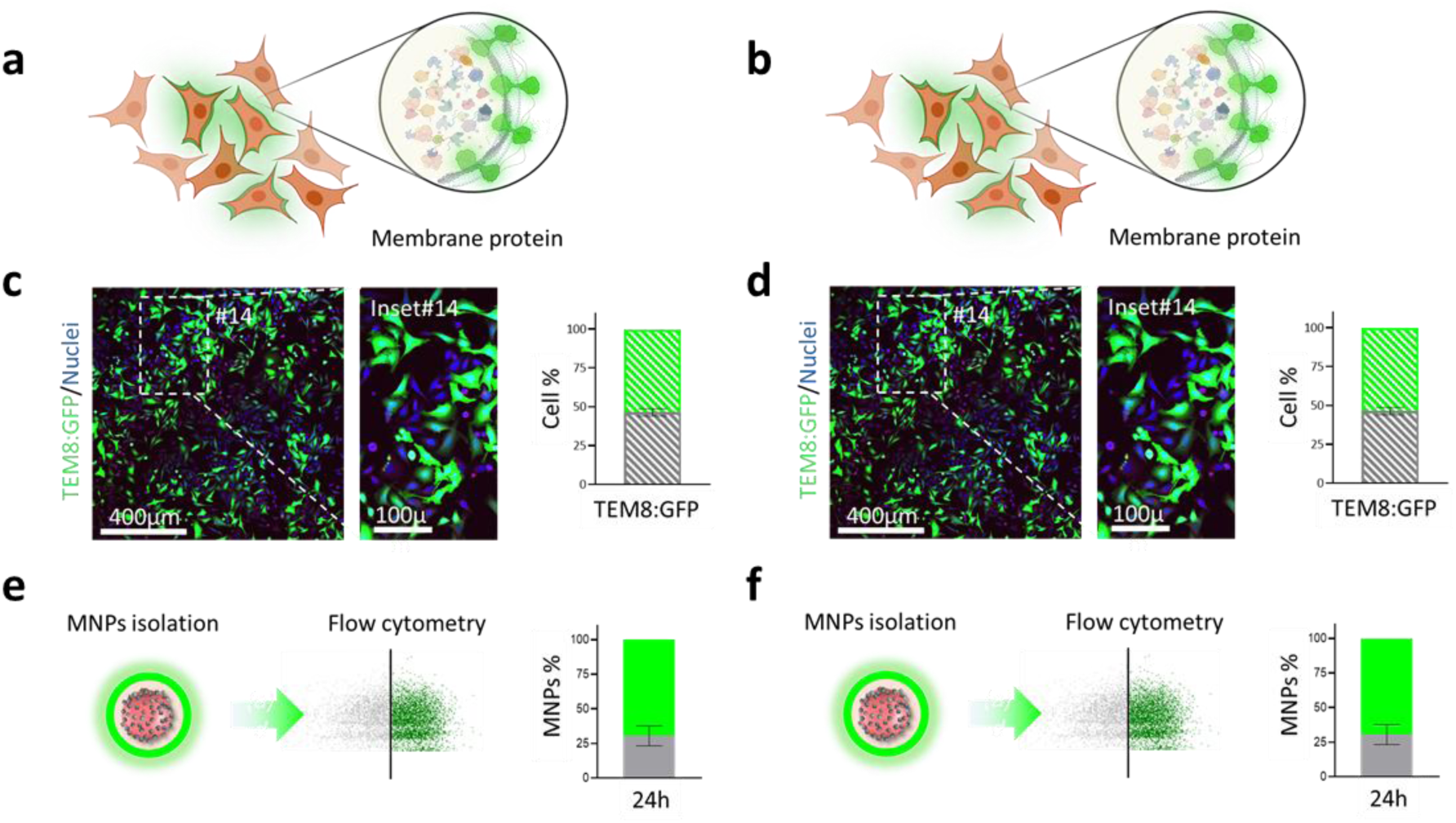
Validation of population-level representativeness. (a,b) Schematic overview of the experimental design using parental cell populations expressing either (a) a cytoplasmic protein (Hsp70:GFP) or (b) a membrane-associated protein (TEM8:GFP). (c, d) Representative confocal images and flow cytometry histograms of the parental populations, showing the ratio of GFP-positive (green) to GFP-negative (blue) cells. (e, f) Flow cytometry analysis of nanobiopsy-derived MNPs. Representative dot plots and accompanying histograms illustrate the fraction of retrieved MNPs carrying (e) Hsp70:GFP or (f) TEM8:GFP signals. The high concordance between the parental cell distributions and the MNP-derived signals demonstrates that magnetic nanobiopsy quantitatively reports cellular heterogeneity. Data are presented as mean ± SD (*n* = 3 for each condition).

Following the 24 h sampling period, the proportions of GFP-positive MNPs were found to broadly align with the distributions characterized in the parental populations. In the case of cytoplasmic sampling, 68 ± 8 % of the parental cells expressed Hsp70:GFP, while 67 ± 2 % of the recovered MNPs were positive, demonstrating an exceptional correlation for soluble cargo. For the membrane-associated TEM8:GFP, 54 ± 2 % of the cells were positive compared to 69 ± 7 % of the recovered MNPs. In both cases, the proportion of samples in the extracted fractions that displayed the fluorescent proteins was statistically equivalent to the parental populations, with no significant differences observed for the cytoplasmic protein (Hsp70:GFP, *p* = 0.90, *df* = 4), or the membrane protein (TEM8:GFP, *p* = 0.40, *df* = 4), confirming the robustness of the sampling method and the platform’s utility as a representative proxy for monitoring population-level trends across different cellular compartments.

Together, these findings demonstrate that magnetic nanobiopsy provides a powerful, non-destructive alternative to bulk lysis for the molecular phenotyping of heterogeneous populations. The successful retrieval of membrane-integrated proteins further validates the budding-mediated extraction mechanism, proving that the MNPs effectively recruit and carry lipid bilayer components during their exit. By generating a pool of probes that broadly aligns with the parental cell composition, this platform offers a unique strategy for longitudinal studies, enabling the monitoring of dynamic shifts in subpopulation composition and phenotypic transitions without the need for terminal sampling.

## 4 Conclusion

This study introduces magnetic nanobiopsy as a conceptually unique and non-destructive platform for longitudinal intracellular sampling, overcoming the historical trade-off between molecular access and cell viability. Unlike conventional interrogation methods which are often terminal or technically prohibitive, our approach enables the repeated retrieval of biomolecules while preserving long-term cellular health. This transformative capability allows for the direct observation of temporal molecular trajectories within the same living population, shifting the paradigm from static snapshots to dynamic, real-time interrogation of biological systems.

The fundamental novelty of this platform lies in its magnetically driven transport mechanism. Our results demonstrate that the probes not only enter the cytosol in a biocompatible manner but also recruit a diverse array of intracellular cargo. The extraction process, driven by magnetic field reversal, culminates in a budding-mediated exit where the probes are recovered enveloped by a host-derived membrane. This natural “packaging” is crucial, as it confirms the capture of both cytosolic elements and structurally constrained membrane components, effectively mirroring the molecular landscape of the parental cell without compromising its integrity.

Equally important is the exceptional accessibility of this technology. By relying on magnetic actuation and standard laboratory equipment, magnetic nanobiopsy is a highly cost-effective and scalable method that can be easily adopted by any research facility without the need for specialized microfabrication or expensive instrumentation. Furthermore, the recovered nanoprobes are fully compatible with conventional analytical gold standards, including flow cytometry, confocal microscopy, PCR, and biochemical assays. Given its simplicity and economy, we envision this platform becoming a standardized tool for the next generation of longitudinal studies, providing a versatile window into the adaptive dynamics of living cells.

## Acknowledgments

The authors acknowledge the financial support from the Spanish Instituto de Salud Carlos III, under Project ref. DTS24/00023, PI22/00030 and PI23/00261 co-funded by the European Regional Development Fund, “Investing in your future; the Gobierno Regional de Cantabria and IDIVAL for the project Refs INNVAL25/11, DTEC24/01: DTEC25/02. The authors gratefully acknowledge Ms. Claudia Zamarron for their technical help, The figures and graphs have been created with BioRender software (BioRender.com, License ID: 9519A1C8-0002).

## Authors’ Contributions

D.M.G., L.G.H. and M. L.F. performed the experiments. All authors discussed the results and wrote the manuscript. L.G.H. and M.L.F. obtained the funding. All authors have approved the final version of the manuscript.

## SUPPLEMENTARY MATERIAL

**Fig. S1.**
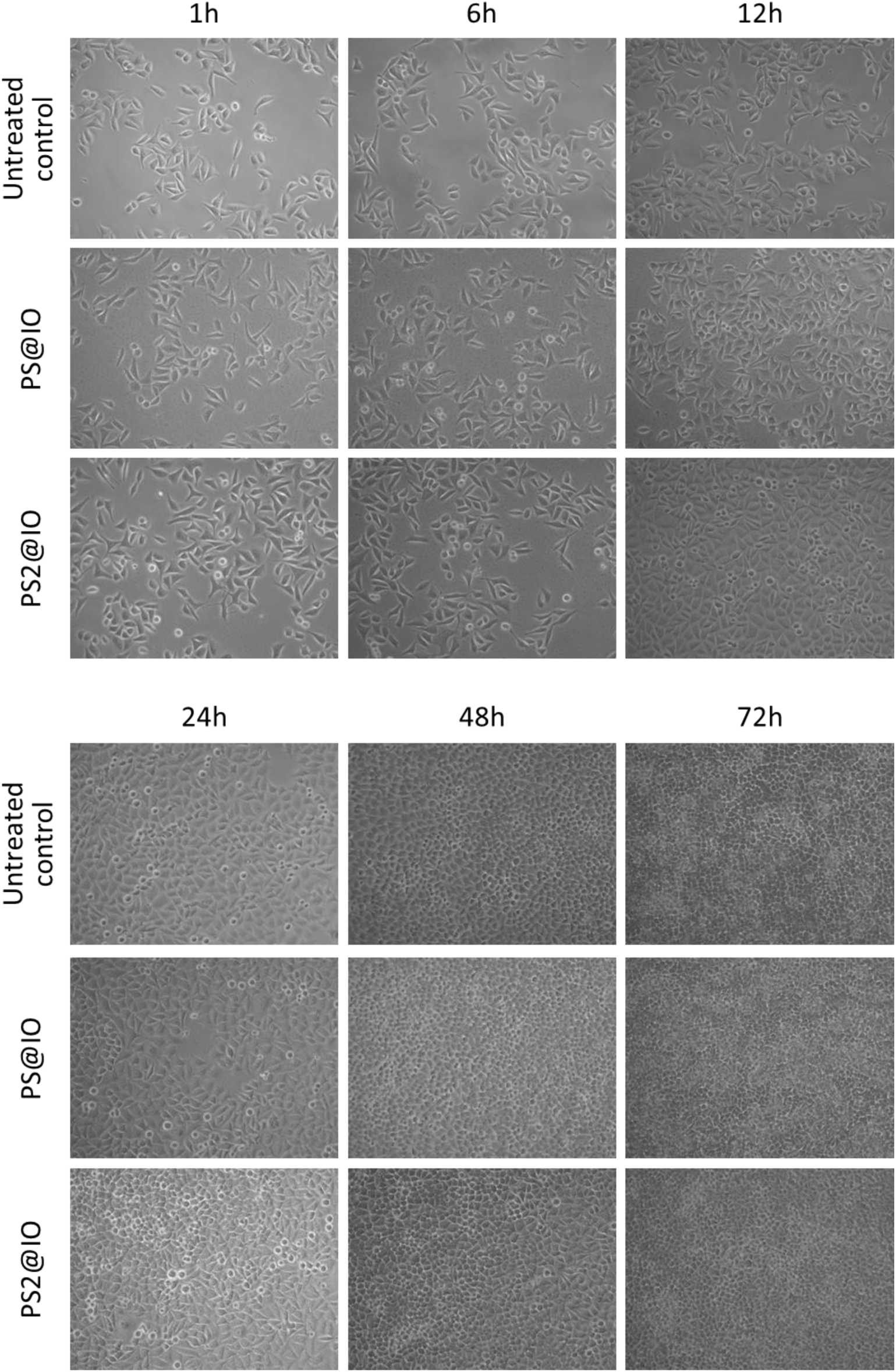
Longitudinal assessment of cell viability via phase-contrast microscopy. Representative images of control cultures and those treated with the indicated MNPs at key experimental time points. The observed cell morphology remains consistent across groups, with no detectable signs of necrosis, apoptosis, or detachment, confirming the maintenance of cellular homeostasis throughout the protocol.

**Fig. S2.**
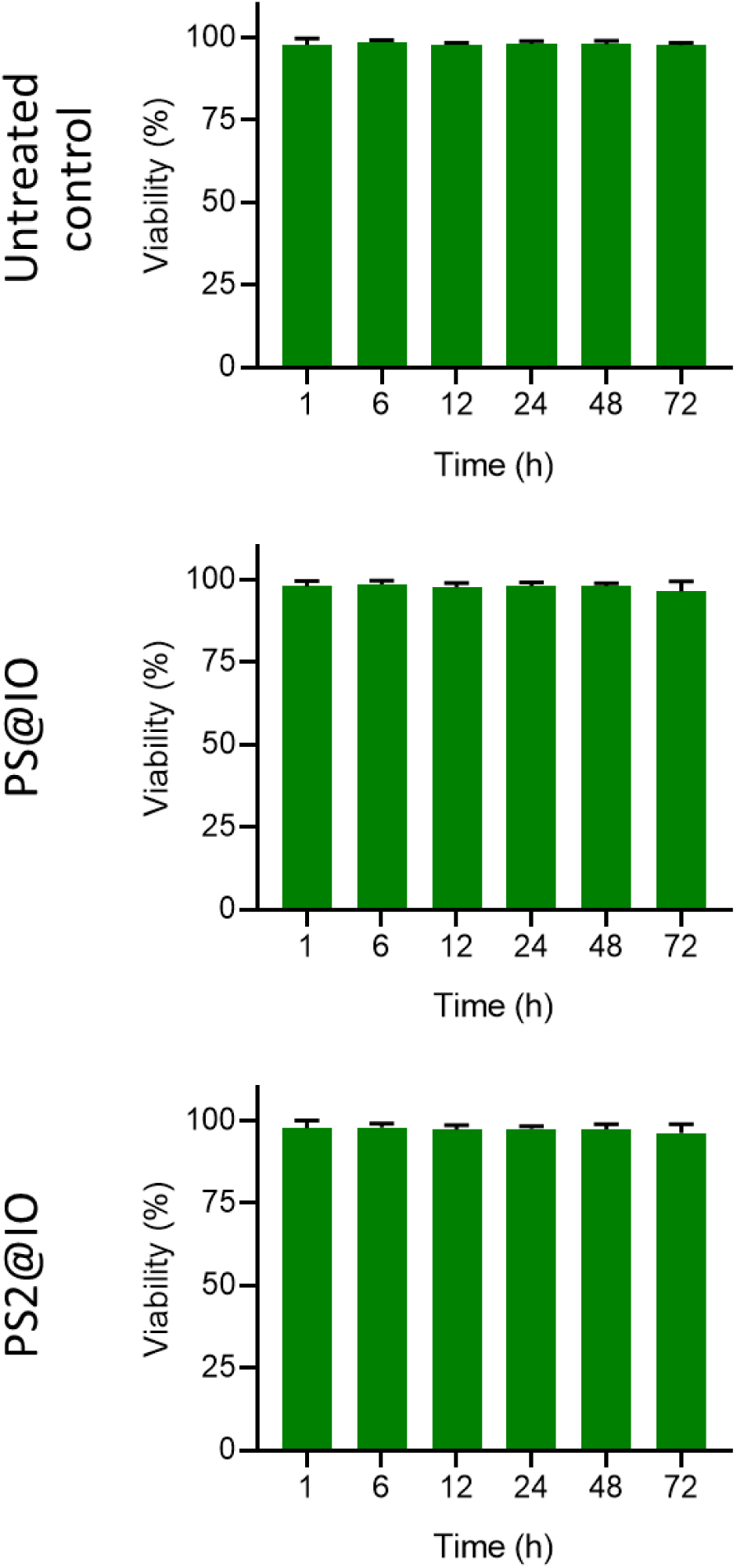
Quantitative analysis of cell viability across experimental stages. Comparative survival rates at discrete time points for: (i) control cultures exposed to the magnetic field without MNPs, and (ii) cultures treated with the indicated MNPs under magnetic actuation. Data represent the mean ± SD, confirming that neither the magnetic field nor the particle-mediated processes significantly impact long-term cell viability.

**Fig. S3.**
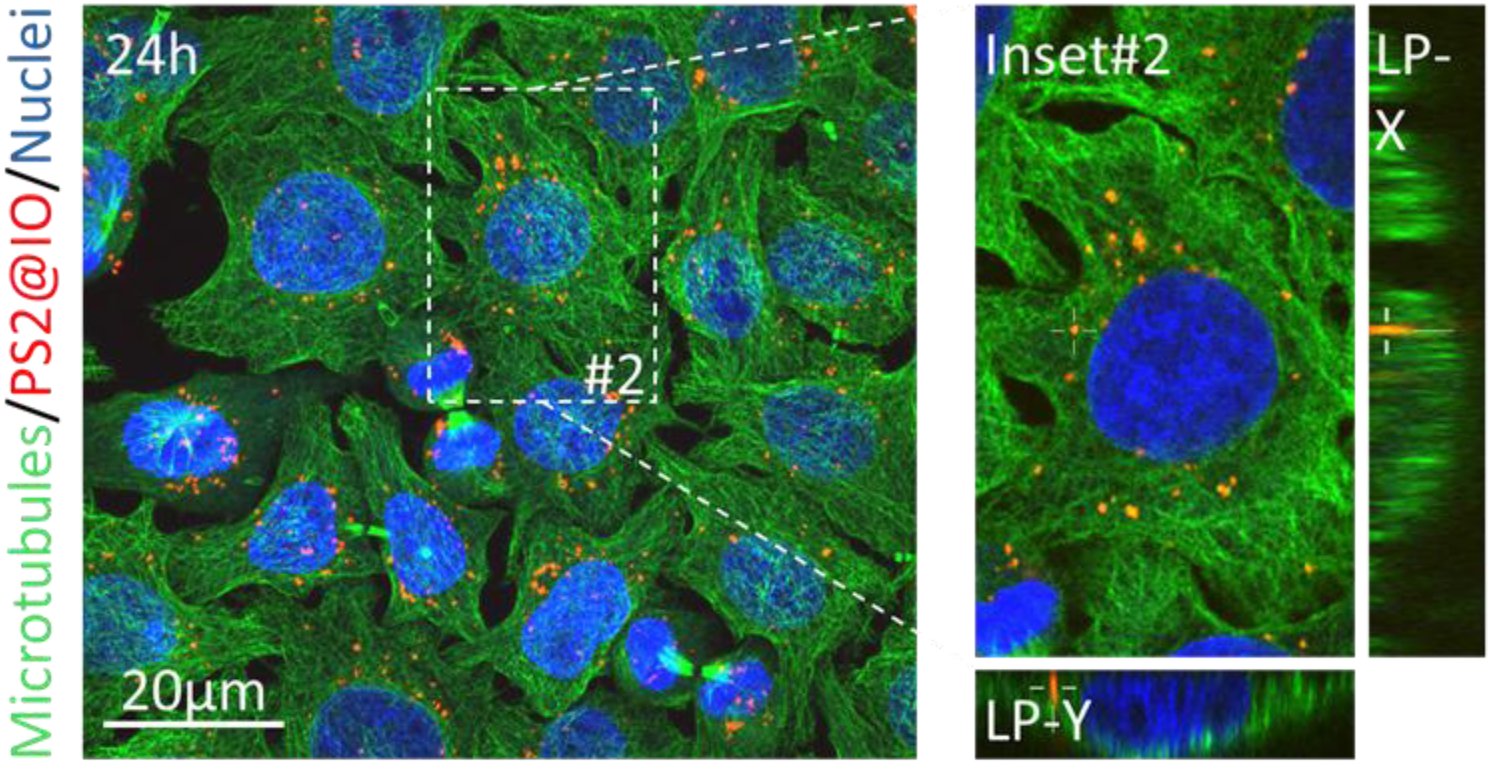
Assessment of cytoskeletal integrity and PS2@IO localization. Confocal microscopy (maximum intensity projections) of cells after 24 h of incubation with PS2@IO nanocomposites (red). The microtubule network was visualized via tubulin immunostaining (green). Inset 1 provides an orthogonal (XZ) lateral projection, confirming the genuine intracellular internalization of the MNPs. The preservation of the microtubule architecture demonstrates that the cytoplasmic residence of the nanoprobes does not induce structural cytoskeletal stress.

**Fig. S4.**
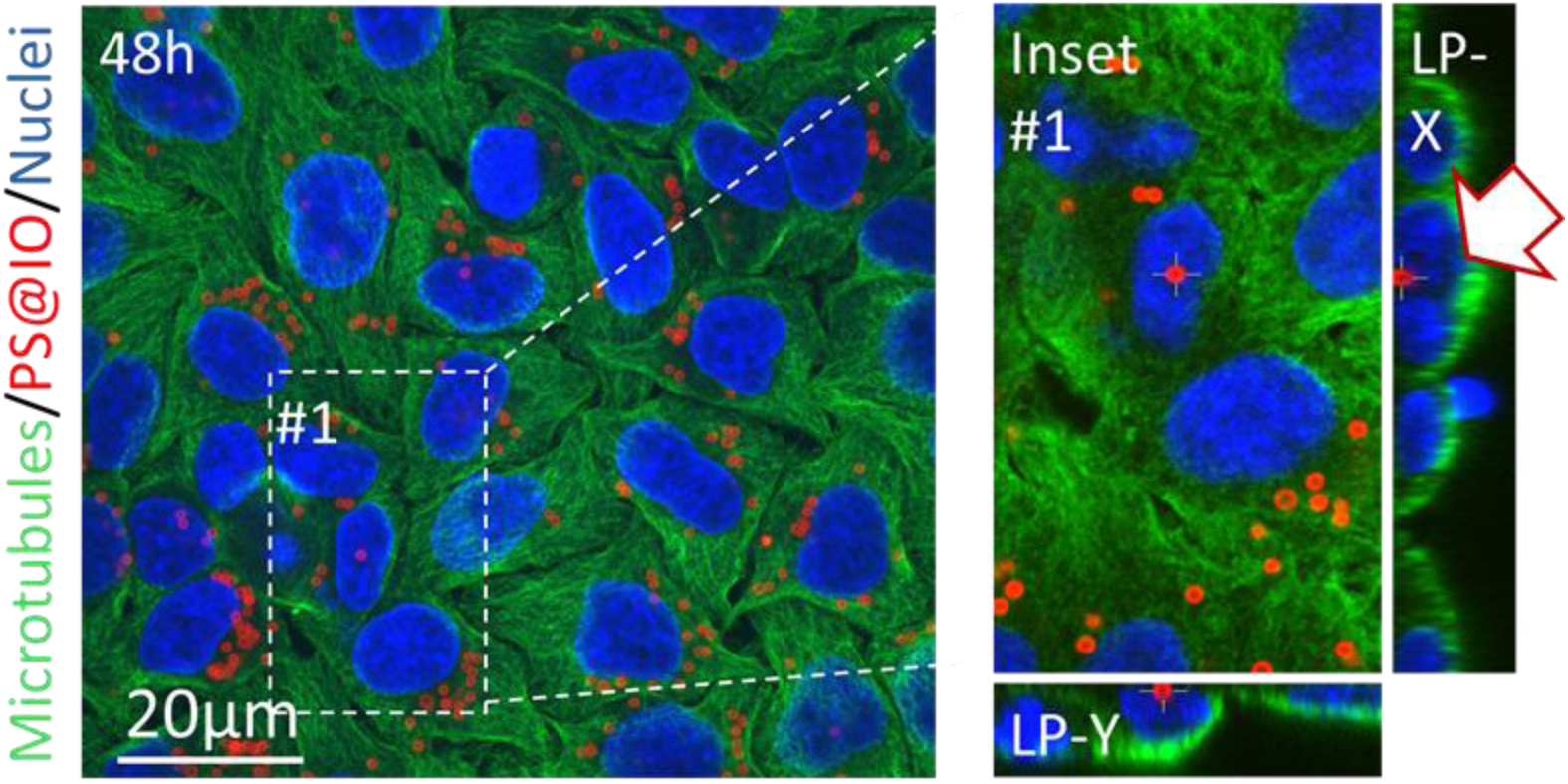
Assessment of cytoskeletal integrity and PS@IO localization at 48h. Confocal microscopy (maximum intensity projections) of cells cultured for 48 h following treatment with the MNP (red). The microtubule network was visualized via tubulin immunostaining (green). Inset 1 displays an orthogonal (XZ) projection, confirming the genuine intracellular localization of the MNPs. The arrow indicates a nanoprobe localized within the nuclear compartment, as verified by the cross-sectional lateral views. The continuous and well-defined tubulin network demonstrates the absence of structural cytoskeletal disruption.

**Fig. S5.**
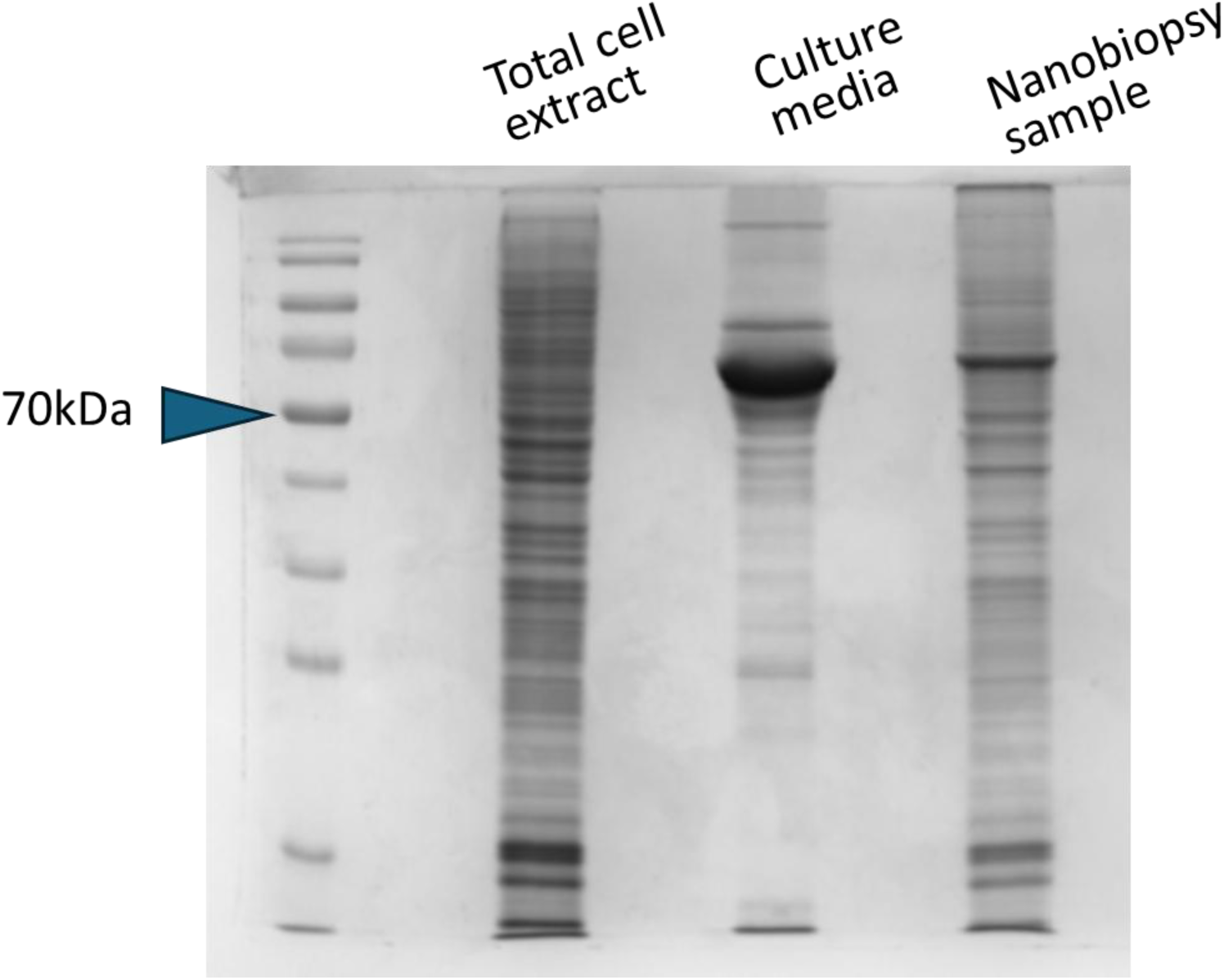
Comparative protein profiling via SDS-PAGE. Representative Coomassie strained electrophoresis gel illustrating the protein landscapes of the parental cell lysate, the culture medium, and the extracted, pooled MNPs harvested from the same population. The distinct banding pattern of the MNP fraction demonstrates the successful enrichment of intracellular proteins and validates the selectivity of the nanobiopsy process against extracellular contaminants.

**Fig. S6.**
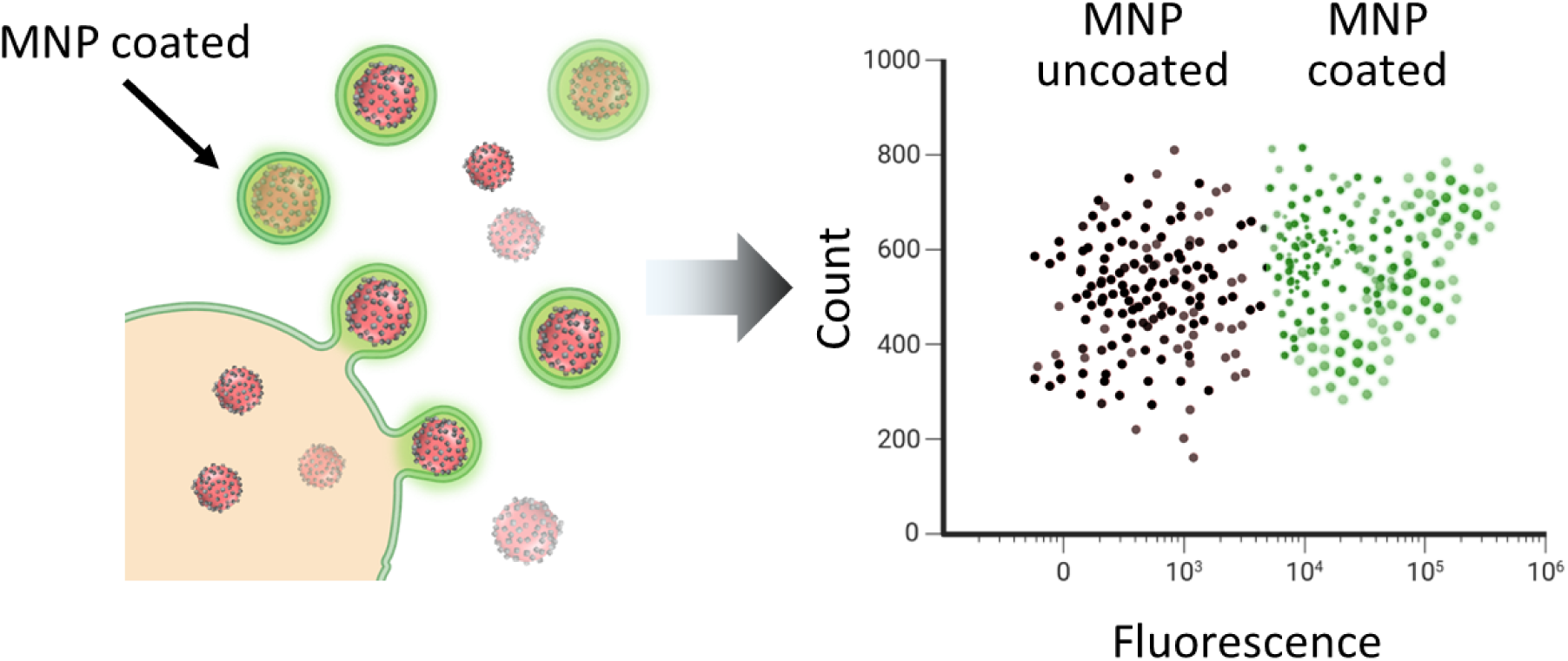
Schematic of the proof-of-concept experimental workflow using extracted MNPs to quantify the proportion of probes capturing cytoplasmic or membrane-associated proteins. The diagram illustrates how the nanobiopsy-derived MNPs reflect the relative abundance of protein-positive and -negative cells in the parental population.

